# Recombinant Human Proteoglycan 4 (rhPRG4) Downregulates TNFα-Stimulated NFκB Activity and FAT10 Expression in Human Corneal Epithelial Cells

**DOI:** 10.1101/2022.10.12.511806

**Authors:** Nikhil G Menon, Yasir Suhail, Ruchi Goyal, Wenqiang Du, Adam P Tanguay, Gregory D. Jay, Mallika Ghosh, Kshitiz, Tannin A Schmidt

## Abstract

Dry Eye Disease (DED) is a complex pathology affecting millions of people with significant impact on quality of life. Corneal inflammation, including via the NFκB pathway, plays a key etiological role in DED. Recombinant human proteoglycan 4 (rhPRG4) has been shown to be a clinically effective treatment for DED that has anti-inflammatory effects in corneal epithelial cells, but the underlying mechanism is still not understood. Our goal was to understand if rhPRG4 affects TNFα-stimulated inflammatory activity in corneal epithelial cells. We treated hTERT-immortalized corneal epithelial (hTCEpi) cells ±TNFα ±rhPRG4 and performed Western blotting on cell lysate and RNA sequencing. Bioinformatics analysis revealed that rhPRG4 had a significant effect on TNFα-mediated inflammation with potential effects on matricellular homeostasis. rhPRG4 reduced activation of key inflammatory pathways and decreased expression of transcripts for key inflammatory cytokines, interferons, interleukins, and transcription factors. TNFα treatment significantly increased phosphorylation and nuclear translocation of p65, and rhPRG4 significantly reduced both these effects. RNA sequencing identified FAT10, which has not been studied in the context of DED, as a key pro-inflammatory transcript increased by TNFα and decreased by rhPRG4. These results were confirmed at the protein level. In summary, rhPRG4 is able to downregulate NFκB activity in hTCEpi cells, suggesting a potential biological mechanism by which it may act as a therapeutic for DED.

## 1. Introduction

According to the TFOS DEWS II report, dry eye disease (DED) is a “multi-factorial disease of the ocular surface characterized by a loss of homeostasis of the tear film, and accompanied by ocular symptoms, in which tear film instability and hyperosmolarity, ocular surface inflammation and damage, and neurosensory abnormalities play etiological roles” [1]. DED is prevalent in a variety of populations, affecting 10-20% of those between 20 and 40 years of age and more than 30% of those above 70 years of age [2]. Common symptoms of DED include ocular itching and burning, light sensitivity, and visual disturbances [3], and these symptoms significantly affect DED patients’ quality of life. Ocular surface inflammation in particular is often a focus of DED treatments. Many studies have shown elevated levels of inflammatory cytokines in the tears of patients with DED, including IL-1β, IL-6, TNFα, and RANTES [4,5]. A key pathway involved in the production of these cytokines is the Nuclear Factor kappa B (NFκB) pathway [6]. The NFκB protein is most commonly composed of two subunits, a 50 kDa subunit (p50) and a 65 kDa subunit (p65). In a normal state, both subunits are bound to an inhibitor, preventing translocation to the nucleus [7]. During inflammation, the inhibitor is phosphorylated and degraded, allowing the complex to translocate to the nucleus and affect gene expression [7,8]. Additionally, the p65 subunit may itself be phosphorylated, enhancing its transcriptional activity [7]. While there are treatment options for DED available, they are not always effective, often not fully addressing the signs and symptoms of DED [9]. As such, there remains a need for an effective treatment to ease the burden on DED patients.

Proteoglycan 4 (PRG4), also known as lubricin, is a mucin-like glycoprotein classically known for its cartilage lubricating properties [10–13]. At the ocular surface PRG4 acts as a boundary lubricant [14], and lack of PRG4 results in increased ocular surface damage in mice [15,16]. Full length recombinant human PRG4 (rhPRG4) demonstrates similar lubricating properties to native PRG4 on articular cartilage [17] and on the ocular surface [15]. In a clinical trial (NCT02507934), rhPRG4 topical application was shown to be clinically effective in improving signs and symptoms in DED patients [18]. The safety and efficacy of 150 μg/mL rhPRG4 was compared to 0.18% sodium hyaluronate eye drops in 39 subjects with moderate DED. The rhPRG4 treatment had significant improvements compared to the eye drops in both symptomatic and objective signs, and there were also no observed treatment-related adverse effects [18]. More recent studies have explored rhPRG4’s anti-inflammatory properties, showing that rhPRG4 can bind to and inhibit toll-like receptors as well as decrease nuclear NFκB translocation in synovial fibroblasts [19,20]. rhPRG4 has also been shown to have anti-inflammatory properties in the ocular environment [21]. Specifically, rhPRG4 has been shown to reduce inflammation-induced cytokine secretion in human corneal epithelial cells, including RANTES, IP-10, and ENA78 [21]. However, the effects of rhPRG4 on inflammatory pathway activation, including NFκB, in these cells have not been explored.

Motivated by our previous work, we sought to examine if rhPRG4 can directly affect NFκB activation in hTCEpi cells stimulated with TNFα. RNA sequencing revealed a broad anti-inflammatory effect of rhPRG4 administration, with reversal of several key inflammatory pathways upregulated by TNFα stimulation. Transcripts of several genes encoding inflammatory cytokines, in particular interferons, were significantly reversed in TNFα-treated cells after addition of rhPRG4. In addition, rhPRG4 significantly reduced TNFα-stimulated phosphorylation and nuclear translocation of NFκB. RNA sequencing analysis identified a molecule not previously studied in DED, the ubiquitin-like modifier human leukocyte antigen-F adjacent transcript 10 (FAT10). FAT10 has been shown to play a proinflammatory role in a variety of organ systems, especially in the context of cancers, and to specifically upregulate TNFα-stimulated NFκB activation. rhPRG4reduces FAT10 expression, which may play a role in its ability to modulate NFκB activity.

## 2. Results

### 2.1. rhPRG4 reduces TNFα-mediated inflammation in hTCEpi cells

We treated hTCEpi cells with rhPRG4 and/or TNFα and collected and analyzed RNA by sequencing analysis to systematically explore rhPRG4’s anti-inflammatory effect (Figure 1a). Principal Component Analysis (PCA) presented a picture indicating that the effect of rhPRG4 was overall independent of inflammatory challenge. While TNFα had a profound effect on hTCEpi cells along the PC1 axis, the effect of rhPRG4 was in the PC2 axis, both without or with TNFα stimulation (Fig 1b). Unbiased analysis of gene sets activated by TNFα stimulation expectedly showed large and significant activation of numerous inflammatory pathways, including response to interferon gamma, type I interferon, chemokine activity, antigen presentation, adaptive immune response, and host response to virus. Interestingly, we found that TNFα stimulation also resulted in significant increase in gene sets associated with extracellular matrix production as well as biogenesis of plasma membrane and endoplasmic reticulum. Treatment of rhPRG4 resulted in significant reversal in activation of these gene sets, particularly those related to inflammation (Fig 1c). We then analyzed the ingenuity pathway analysis (IPA) predicted activation and de-activation of factors by TNFα and rhPRG4 treatment, singly and in combination (Fig 1d). The results showed that the effect of rhPRG4, in combination with TNFα stimulation, was specific for inflammation related factors. rhPRG4 reversed TNFα activation of *IFNA2* (interferon α), *IFNB1* (interferon β), *IFNG* (interferon γ), *IL1A* (interleukin-1α), and *IL1B* (interleukin-1β), as well as *OSM* (oncostatin), which is involved in IL2 activation. In contrast, no coherent biological signal could be interpreted from predicted activation of factors that changed with rhPRG4 in parallel to TNFα, or those which had decreased in TNFα treatment but increased with rhPRG4. These patterns suggested that rhPRG4 treatment has a specific anti-inflammatory effect on cell challenged with an inflammatory stimulus, namely TNFα.

**Figure 1.**
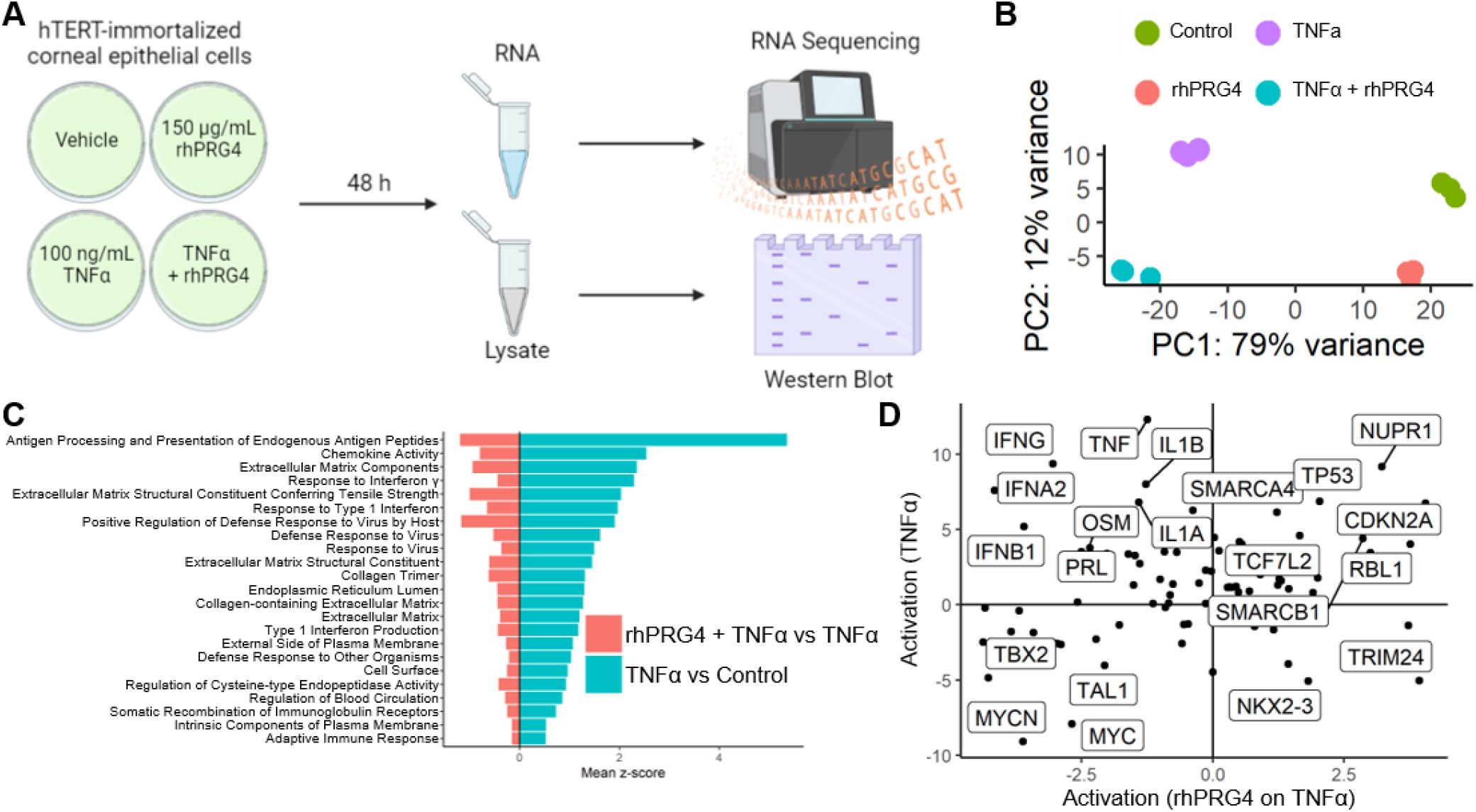
The effect of rhPRG4 on TNFα-stimulated inflammatory activity at the RNA level (A) Experimental setup for collecting RNA and cell lysate from hTCEpi cells. Created with BioRender.com (B) Principal component analysis of the expression of 500 most variable genes showing the consistent large (~ 79% variance) effect of the TNFα and the effect of rhPRG4 on the x and y-axes. (C) Gene Ontology gene sets that are positively regulated by TNFα on control and down-regulated by the application of rhPRG4 on TNFα treated samples. (D) The activation z-score (calculated using the expression of downstream genes) of upstream regulators for TNFα treatment (y-axis) and the addition of rhPRG4 treatment on TNFα (x-axis).

### 2.2. rhPRG4 reduces TNFα-stimulated p65 phosphorylation in hTCEpi cells

Focusing on activated upstream regulators which increased in response to TNFα treatment but showed significant reversal after rhPRG4 treatment, we again found the set populated by genes encoding for key modulators and actors in inflammation, such as genes encoding for interferons (*IFNA2, IFNB1*) and interleukins (*IL1A, IL1B, IL17A, IL15, IL2, IL3*, and *IL6*). Additional genes of interest were *RELA*, which encodes for p65, a component of the NFκB complex that acts as a key inflammation-related transcriptional activator in response to TNFα stimulation, and *SP1,* which transcriptionally regulates many inflammatory genes including CXCL4 [22], and ligand for CD40 (*CD40LG*), which costimulates inflammatory response in many systems including in the vasculature (Fig 2a) [23]. We therefore sought to experimentally confirm if rhPRG4 treatment indeed directly reduced p65 signaling, the chief effector of a TNFα-mediated inflammatory response.

**Figure 2.**
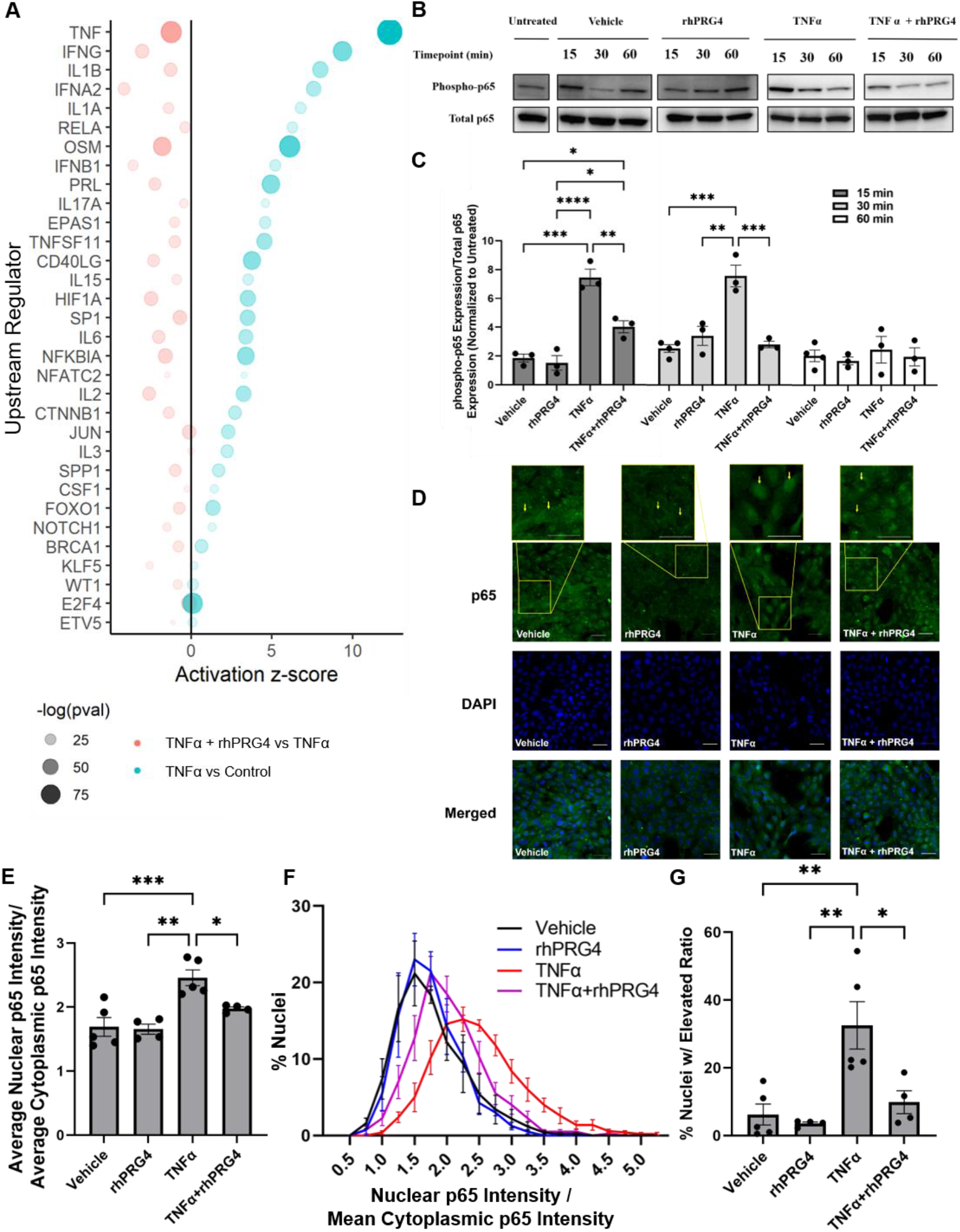
The effect of rhPRG4 on upstream regulators stimulated by TNFα, specifically NFκB. (A) Upstream regulators that were activated by TNFα treatment, and down-regulated thereafter on the addition of rhPRG4. Size and shading correspond to the statistical significance (p-value) for the enrichment of downstream regulation while the x-axis is the overall effect size of the activation. (B) p65 phosphorylation is stimulated by TNFα and downregulated by rhPRG4. p65 phosphorylation analyzed by Western Blotting (N=3-4) using anti-phospho-p65 and anti-p65 antibodies. (C) Relative expression quantified using Genetools and normalized to expression level of untreated sample. (D) Internalization of p65 after 30 min analyzed by immunofluorescence staining with an anti-p65 antibody. (E) Relative mean nuclear and cytoplasmic p65 intensities quantified with ImageJ, and the ratio of nuclear to cytoplasmic intensity calculated (N=4-5). (F) Ratio of nuclear to cytoplasmic p65 intensities calculated for each nucleus, shown as histogram (N=250-553). (G) Percentage of nuclei with nuclear to cytoplasmic ratios 2 standard deviations above vehicle alone (N=4-5).

A time-course Western blot analysis of hTCEpi cells after stimulation with TNFα showed a sharp spike in phosphorylated p65 abundance, which has improved transcription activity, after 15 minutes (Figure 2c, p<0.001). This then stabilized at a plateaued level comparable to vehicle treatment after 60 minutes. The addition of rhPRG4 alone did not significantly influence phosphorylated p65 expression at any timepoint. However, the addition of rhPRG4 to TNFα-stimulated hTCEpi cells significantly reduced phosphorylated p65 expression compared to TNFα alone after 15 minutes (Figure 2c, p<0.01) and 30 minutes (Figure 2c, p<0.001).

Subsequent to phosphorylation, p65 translocates to the nucleus acting as key transcription factor jumpstarting many inflammation-related genes [7,8]. Immunofluorescence of hTCEpi cells confirmed our earlier findings of a robust p65 accumulation in response to TNFα treatment which was significantly reversed with rhPRG4 treatment. At the 30-minute timepoint, rhPRG4 alone did not significantly affect nuclear NFκB expression (Figure 2d,e), TNFα significantly stimulated nuclear NFκB levels (Figure 2d,e, p<0.001), and rhPRG4 was able to reduce nuclear NFκB expression after stimulation (Figure 2d,e, p<0.05). When nuclei are ordered by their nuclear to cytoplasmic p65 ratio, and a cutoff is selected where the majority of the nuclei that received only the vehicle are below the cutoff (Figure 2f), TNFα treatment significantly increases the percentage nuclei above the cutoff (Figure 2g, p<0.001), and rhPRG4 was able to significantly reduce the percentage of nuclei with an elevated nuclear to cytoplasmic p65 ratio (Figure 2g, p = 0.05).

### 2.3 Identifying TNFα-activated genes specifically reversed in expression by rhPRG4

Our data strongly indicated that rhPRG4 administration has a broad anti-inflammatory effect in TNFα-challenged hTCEpi cells. To understand plausible molecular effectors of this response, we sought to identify genes whose gene expression was specifically reduced by rhPRG4. Gene set enrichment analysis (GSEA) [24], a non-parametric approach to interpret involvement of gene-sets in the comparative data, showed high positive enrichment of NFκB downstream genes in response to TNFα stimulation and rhPRG4 treatment reversed NFκB enrichment (Figure 3). We then identified genes which constituted the leading edges of NFκB positive enrichment in a comparison between TNFα-treated and control and the leading-edge genes of NFκB negative enrichment in a comparison between TNFα+rhPRG4 and TNFα conditions. The leading edges indicate the genes most responsible for the enrichment of the given gene-set. The intersection of these leading edges revealed a gene-set which was reversed in expression specifically by rhPRG4 (Figure 3). These genes included several inflammation-associated C-X-C motif chemokines (*CXCL1/5/9/10/11*) and C-C motif chemokine ligands (*CCL2/5*), several of which were previously shown to be downregulated by rhPRG4 at the protein level [21], various adhesion receptors, including *ITGAV* and *ITGAM*, which encode for alpha subunits of integrins, and *VCAM1* and *NCAM1*, which encode for adhesion receptors of the immunoglobulin family. Of particular note in this gene-set was the gene *UBD,* which encodes for a ubiquitin-like molecule, FAT10. FAT10 has been found to be highly upregulated in a variety of cancers [25] and, importantly in this study, to be an upregulator of NFκB activation [26].

**Figure 3.**
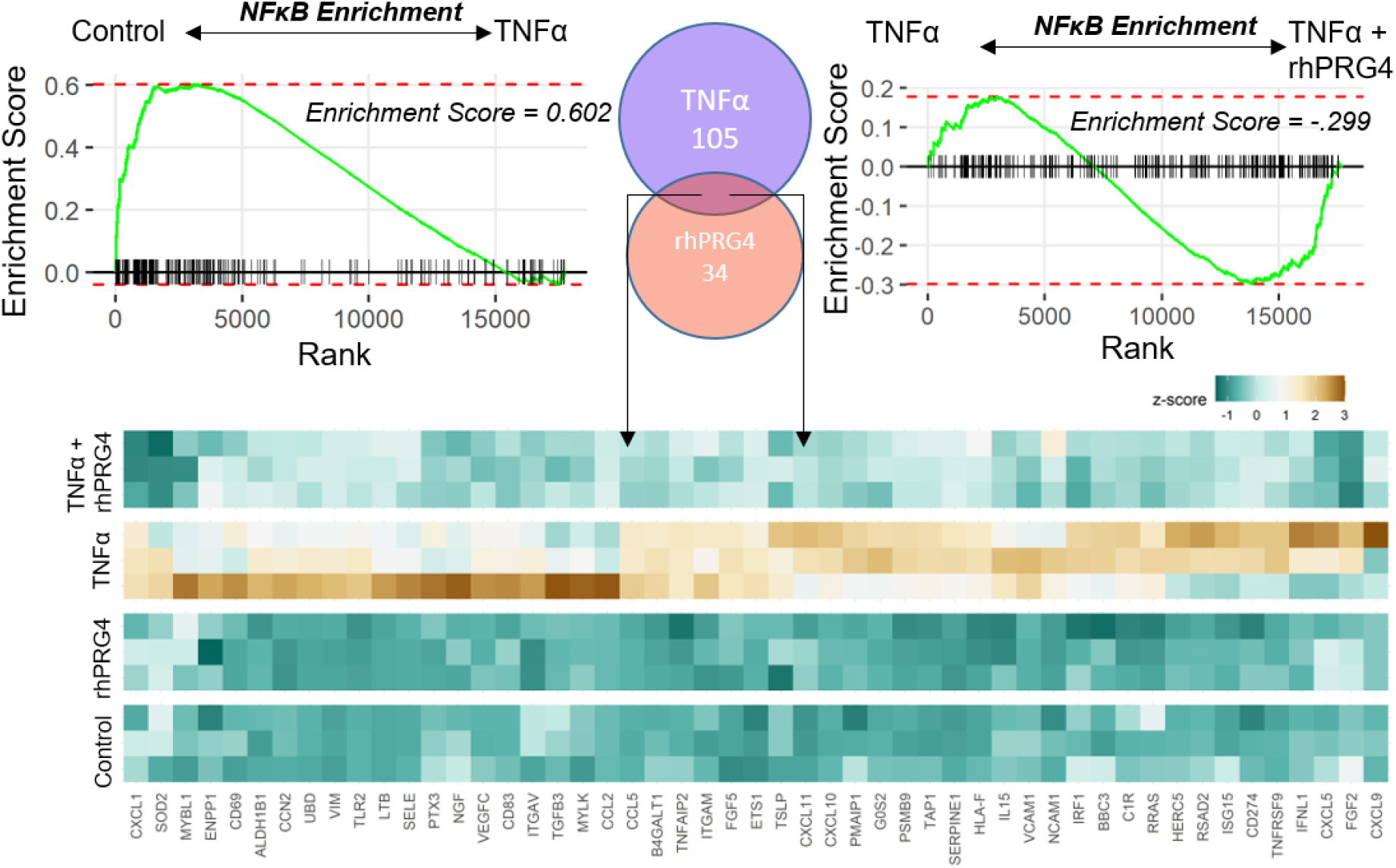
Gene Set Enrichment Analysis diagram shows the enrichment of NFκB complex genes in the up-regulated genes on TNFα treatment and the relative enrichment in down-regulated genes upon rhPRG4 application on TNFα treatment. The Venn diagram shows the number of NFκB genes exclusively highly down-regulated by rhPRG4 and up-regulated by TNFα, according to the GSEA leading edges. The common set of genes in the intersection are represented in the heatmap showing the regulation by TNFα and rhPRG4.

### 2.4. rhPRG4 reduces TNFα-stimulated FAT10 RNA and protein expression

We identified genes that were modulated by rhPRG4 specifically within the set of TNFα-stimulated genes. Among genes identified, an interesting candidate discovered was *UBD*, encoding for FAT10 protein, which has been previously shown to regulate IκB stability and to be upregulated itself by NFκB (Figure 4a). IκB, when ubiquitinated, is targeted for proteasomal degradation, releasing p65 to translocate to the nucleus and start transcription. Direct ranking of genes significantly regulated by either TNFα or rhPRG4, singly and in combination, also featured UBD (Fig 4b). We therefore tested if FAT10 abundance was regulated by rhPRG4. Examining RNA expression (reads per million reads, RPM) of FAT10 specifically revealed that TNFα significantly stimulated FAT10 RNA expression (Figure 4c, p<0.01), and rhPRG4 addition significantly decreased this stimulated expression (Figure 4c, p<0.05). TNFα also significantly stimulated FAT10 protein expression (Figure 4e, p<0.01), and rhPRG4 significantly reduced FAT10 protein expression with TNFα stimulation (Figure 4e, p<0.0001) while not affecting FAT10 protein expression in the absence of TNFα.

**Figure 4.**
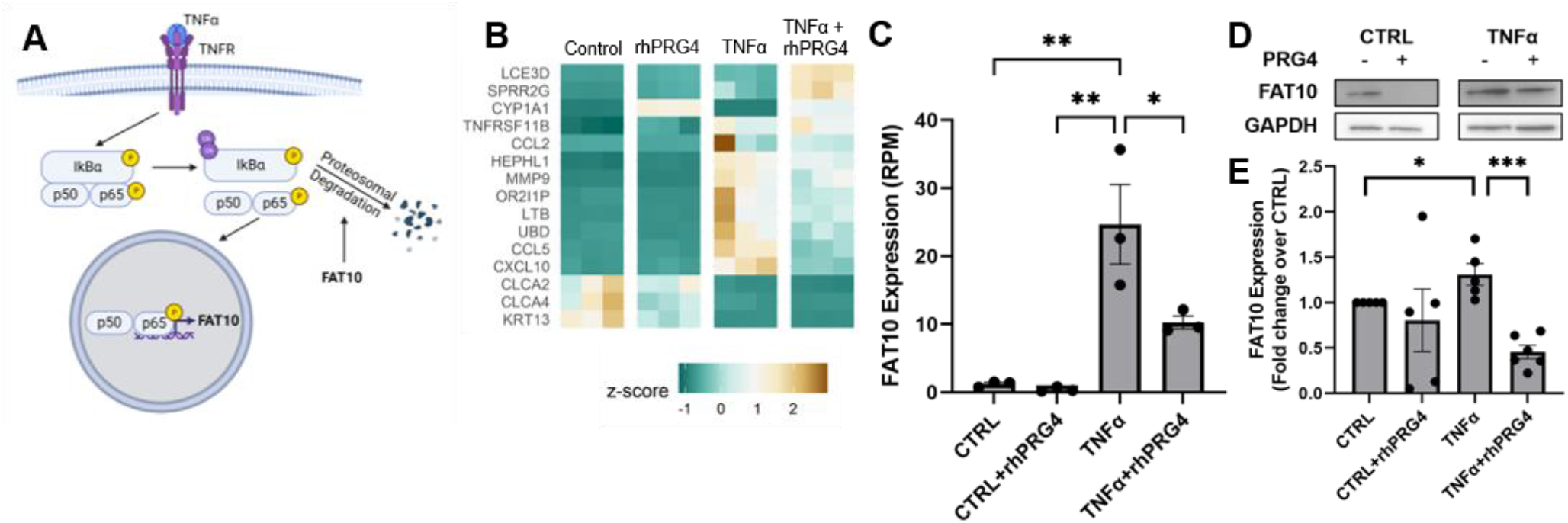
The effect of rhPRG4 on FAT10 expression. (A) FAT10’s role in NFκB signaling. Created with BioRender.com (B) Heatmap showing genes that were most significantly changed by TNFα and rhPRG4. (C) FAT10 RNA expression in RPM from RNA sequencing (C, N=3). (D) FAT10 expression on the protein level by Western Blotting. (E) Relative FAT10 expression normalized to GAPDH expression and to CTRL samples, analyzed using GeneTools software (N=5-7)

## 3. Discussion

This work represents a systems understanding of the effect of rhPRG4 on cells subjected to inflammatory stimuli. We show that treatment of human corneal epithelial cells with rhPRG4 can dramatically reverse TNFα-driven inflammation, which plays a key role in DED. We showed that rhPRG4 reduced both p65 phosphorylation and nuclear translocation after stimulation with TNFα and also identified a key signaling molecule, FAT10, which is stimulated by TNFα and can be downregulated by rhPRG4. This protein has not been studied previously in the context of DED, though it often mediates inflammation in other diseases and organ systems, and it could represent a novel target for therapeutics and potential mechanism through which rhPRG4 improves signs and symptoms of DED.

The results from RNA sequencing represent novel insights into the mechanisms by which rhPRG4 acts to downregulate inflammation induced by TNFα. The increase in the gene expression of various cytokines and chemokines, including RANTES, IP-10, and MCP-1, had previously been observed at the protein level [21] thus validating the findings here. An overlap gene set enrichment analysis identified potentially NFκB pathway related genes which specifically reverse their TNFα-induced expression in response to rhPRG4, suggesting new areas of study to better understand the mechanisms of rhPRG4’s anti-inflammatory effects. VCAM1, for example, plays a key role in leukocyte trafficking, and its binding partner, VLA-4, has been shown to be an effective target for numerous inflammatory diseases and specifically for DED [27].

Phosphorylation of p65, one of the subunits of NFκB, can modulate NFκB activity through the classical pathway and contribute to another IκBα-independent pathway to NFκB translocation. Previously, it has been shown that p65 can be phosphorylated in response to TNFα, and that phospho-p65 has a lower binding affinity to IκBα [28,29]. Additionally, phospho-p65 can translocate to the nucleus regardless of the presence of the proteosome inhibitor MG-132, which blocks the degradation of IκBα [29]. NFκB activity through the classical pathway can also be enhanced by p65 phosphorylation as phosphorylated p65 exhibits enhanced transcriptional activity [30]. Indeed, the maximal NFκB transcriptional response is only achievable through combined activation of the classical pathway and p65 phosphorylation [7]. In the past, rhPRG4 has been shown to reduce NFκB activity in both synoviocytes [19,31] and TLR-HEK cells [32]. In synoviocytes, this downregulation occurs through inhibition in IκBα phosphorylation [19]. Here, we show another mechanism by which rhPRG4 can control NFκB, namely through decreasing p65 phosphorylation. While rhPRG4’s effect on IκBα phosphorylation in corneal epithelial cells was not studied here, this could be explored in future work, in addition to rhPRG4’s effect on other inflammatory transcription factors, including p38 and IRF3.

RNA sequencing identified FAT10 as a molecule of interest that is highly upregulated by TNFα and subsequently downregulated by rhPRG4. FAT10, also known as diubiquitin, is a ubiquitin-like molecule typically found in immune cells, including mature dendritic cells and B cells [33], but it has also been shown to be synergistically inducible in a variety of cell types by TNFα and IFNγ [34]. FAT10 can conjugate proteins, marking them for degradation in a similar manner to ubiquitin. FAT10, and its related gene *UBD*, has not been extensively studied in the context of DED, but gene expression of *UBD* has been shown to be upregulated in the lacrimal glands of non-obese diabetic mice [35]. This mouse model has been used to study Sjogren’s Syndrome, a disease which can also cause patients to suffer from dry eye FAT10 plays an important role in NFκB activation; NFκB activation and translocation, as well as IκBα degradation, were significantly downregulated in FAT10^−/−^ renal tubular epithelial cells stimulated with TNFα [26]. Here, we show that rhPRG4 downregulates TNFα-stimulated FAT10 expression in human corneal epithelial cells at the protein level, indicating another potential mechanism by which it can modulate NFκB activity. Future work could further examine the role FAT10 plays in DED and how downregulating FAT10 with rhPRG4 directly affects cells in the context of DED-associated inflammation, thereby enhancing our understanding of rhPRG4 as a therapeutic.

## 4. Materials and Methods

### 4.1 Cell culture

hTert-immortalized corneal epithelial (hTCEpi) cells, provided by Dr. James Jester [36], were grown and differentiated, as described previously [21,37]. Briefly, hTCEpi cells were grown in Keratinocyte Serum-Free Medium supplemented with 5 ng/mL epidermal growth factor and 25 μg/mL bovine pituitary extract (Gibco, Grand Island, NY) and differentiated in DMEM/F12 (Gibco) + 10% FBS (R&D Systems, Minneapolis, MN) + 10 ng/mL EGF (Gibco).

#### 4.2.1 Cell Treatment and Sample Collection – Time course for NFκB activity and translocation

hTCEpi cells were plated in 60mm dishes at a density of 1,000,000 cells/dish, as well as in a glass bottom 96-well dish at a density of 2,000 cells/well, and allowed to grow for 2 days. Then, the cells were differentiated as described above for 4 days, with a media change on the third day. Two hours before treatment, all wells were changed to DMEM/F12 + 10 ng/mL EGF. Each dish and well received rhPRG4 (1.33 mg/mL, Lubris Biopharma, Framingham, MA) or sterile PBS containing 0.01% Tween-20 immediately followed by treatment media with or without TNFα (Peprotech, Cranbury, NJ) for 15 min, 30 min, or 60 min. Wells that did not receive any treatments were used as a control. The final concentration of the rhPRG4 was 300 μg/mL, and the final concentration of TNFα was 100 ng/mL in DMEM/F12 + 10 ng/mL EGF. Lysate was collected from the 60mm dishes using a specialized lysis buffer (50 mM Tris-HCl (Invitrogen, Carlsbad, CA), 150 mM sodium chloride (Millipore Sigma, Burlington, MA), 1% NP-40 (Fisher Scientific), 5mM EDTA (Millipore Sigma), 10 μg/mL aproteinin (Millipore Sigma), 10 μg/mL leupeptin (Millipore Sigma), 1 mM sodium fluoride (Millipore Sigma), 1 mM sodium orthovanadate (Millipore Sigma), 1mM PMSF (Millipore Sigma), cOmplete Mini protease cocktail (2x, Millipore Sigma). Lysate samples were analyzed by BCA kit, following manufacturer guidelines, to determine protein concentration. This experiment was repeated three times, for a total of N=3 per group.

#### 4.2.2 Cell Treatment and Sample Collection – Single timepoint for RNA and FAT10 analysis

hTCEpi cells were plated in 12-well plates at a density of 100,000 cells/well and allowed to grow for 2 days. Then, the cells were differentiated as described above for 4 days, with a media change on the third day. Each well received rhPRG4 or PBS + Tween as well as media with or without TNFα as described above for 48h. The final concentration of the rhPRG4 was 300 μg/mL, and the final concentration of TNFα was 100 ng/mL in DMEM/F12 + 10 ng/mL EGF. RNA was collected using GeneJET RNA Purification kit (Thermo Fisher, Grand Island, NY), following manufacturer guidelines. Lysate was collected using Mammalian Protein Extraction Reagent (Fisher Scientific, Grand Island, NY), following manufacturer guidelines. Lysate samples were analyzed by BCA kit (Fisher Scientific), following manufacturer guidelines, to determine protein concentration. This experiment was repeated three times, for a total of N=4-5 per group.

### 4.3 Western Blotting

Sodium dodecyl sulphate-polyacrylamide gel electrophoresis (SDS-PAGE) was performed. For the time course lysate samples, 50 μg of each cell lysate sample was loaded onto a 4-20% Tris-Glycene gel (Invitrogen) and electrophoresed, electroblotted to a PVDF membrane, and blocked in 5% non-fat dry milk (Biorad, Hercules, CA) in Tris-buffered saline + 0.05% Tween-20 (TBST). Membranes were then probed with an anti-phospho-p65 antibody (1:1000, 3033, Cell Signaling Technologies, Danvers, MA) and a goat anti-rabbit secondary antibody (1:3000, SeraCare, Milford, MA) in 5% bovine serum albumin (BSA, Thermo Fisher) in TBST. The membrane was then imaged on a G:Box Chemi XX9 imager (Syngene, Frederick, MD) using ECL Select substrate (Cytiva, Marlborough, MA). Next, the membranes were stripped using Restore Stripping Buffer (Fisher Scientific), blocked in 5% non-fat dry milk in TBST, and reprobed using an anti-p65 rabbit antibody (1:1000, 8242, Cell Signaling Technologies), the same goat anti-rabbit antibody used previously, and ECL Select substrate. Intensities of resulting bands were quantified by densitometry with the Genetools software (Syngene), and the intensity of the 8hosphor-p65 band was normalized to the intensity of the p65 band for each lane.

For the single timepoint lysate samples, 9.5-10 μg of each cell lysate sample was loaded onto a 4-20% Tris-Glycene gel and electrophoresed, electroblotted to a PVDF membrane, and blocked in 5% non-fat dry milk in TBST. Membranes were then probed with a rabbit anti-FAT10 antibody (1:1000, MBS9204261, MyBioSource, San Diego, CA) and a goat anti-rabbit secondary antibody (1:10000, AP307P, Millipore Sigma) in 3% non-fate dry milk in TBST followed by imaging with ECL select substrate as mentioned above. Membranes were then stripped, blocked, and reprobed using a mouse anti-GAPDH antibody (1:1000, MAB374, Millipore Sigma) and a goat anti-mouse secondary antibody (100 ng/mL, A4416, Millipore Sigma), as described above. Membranes were imaged using ECL Select substrate, and intensities of resulting bands were quantified by densitometry with the Genetools software.

### 4.4 RNA sequencing & Bioinformatics

RNA isolated from 3 replicates from each of the 4 treatments (Control, TNFα, rhPRG4, and rhPRG4+TNFα) was sequenced in a paired-end fashion on the Illumina platform. Resulting FASTQ files were aligned to the GRCh38 transcriptome using hisat2. Read pairs mapping concordantly to a unique position on the transcriptome were counted using featureCounts. An average of 11,486,217 reads per sample were counted against known genes, with 17,721 genes having at least one read mapping to them. Differential expression was between conditions was computed using DESeq2 on the R platform. P-values were corrected for multiple testing using the Benjamini-Hochberg method. Effect of the differential expression was measured using log 2-fold change values.

Overall enrichment of differentially expressed genes in gene-sets (Gene Ontology/KEGG) was performed using the following. First, DE p-values were converted to equivalent z-scores using the inverse normal distribution. The sign of the z-score (up- or down-regulation) was taken from the fold change. For each gene-set, the mean of the z-scores in the set vs. all genes outside the set was compared using the Student’s t-test. The size and sign of the enrichment was calculated as the mean z-score of the genes in the set. To find the enrichment and specific genes in the NFκB complex, GSEA was used and the leading-edge genes were counted.

### 4.5 Immunofluorescence

Cells were fixed with 4% paraformaldehyde for 15min, permeabilized with 0.05% Triton-X100 for 10min, and blocked with 5% goat serum and 1% BSA for 1h at room temperature. Cells were then incubated with rabbit NFκB p65 antibody (1:200, 10745-I-AP, Proteintech, Rosement, IL) at 4°C overnight. After washing, cells were incubated with AF488-conjugated goat anti-rabbit secondary antibody (1:400, Invitrogen) for 1h at room temperature. Cell nuclei were counter stained with DAPI. Images were taken using Zeiss Axio Observer Z1 with a 20x objective.

After images were collected, samples were analyzed using ImageJ. The average mean gray value was determined at five locations per sample for nuclear and cytosolic NFκB. Nuclear NFκB was designated as areas with both NFκB and DAPI staining. Cytosolic NFκB was designated as areas within the cell but without DAPI staining. Nuclear NFκB values were normalized to cytosolic NFκB values within each area.

Additional analysis was performed by calculating the mean grey value for each nucleus within each of the five locations for every treatment condition and normalized to the mean cytosolic value for the whole location. A cutoff for separating nuclei by nuclear p65 levels was determined as the mean of the upper limits of the confidence intervals for each vehicle-treated location. The percentage of nuclei above this cutoff was calculated for each location.

### 4.6 Statistical Analysis

Data are expressed as mean ± SEM. The effect of TNFα and rhPRG4 on the phosphorylation of NFκB was assessed by one-way ANOVA with Tukey’s post-hoc testing at each timepoint. The effect of TNFα and rhPRG4 on nuclear translocation of NFκB was assessed by one-way ANOVA with Tukey’s post-hoc testing. The effects of TNFα and rhPRG4 on FAT10 RNA and protein expression were assessed by one-way ANOVA with Tukey’s post-hoc testing. Outliers were identified using the ROUT test with cutoff of 5%

## 5. Conclusions

We show here that rhPRG4 is able to downregulate NFκB activity and translocation in human corneal epithelial cells stimulated with TNFα. We also show, for the first time, that rhPRG4 downregulates TNFα-induced FAT10 expression, a protein that has not been previously studied in the context of dry eye but does play a proinflammatory role in other diseases. Overall, these findings provide a new avenue to study and further understand PRG4’s anti-inflammatory effects, in the context of rhPRG4 as an effective treatment for DED.

## Author Contributions

Conceptualization, M.G., K., and T.S.; methodology, M.G., K., and T.S.; software, Y.S.; validation, N.M., R.G., and A.T.; formal analysis, N.M. and Y.S..; investigation, N.M., R.G., W.D., A.T., and M.G.; resources, T.S., G.J.; data curation, Y.S.; writing—original draft preparation, N.M., K., Y.S., and W.D.; writing—review and editing, T.S., K., G.J., M.G., and A.T.; visualization, Y.S.; supervision, M.G., K., T.S.; project administration, T.S.; funding acquisition, T.S. All authors have read and agreed to the published version of the manuscript.

